# Prevalence of Alcohol use during pregnancy and its association with partner alcohol use in East Africa: systematic review and meta-analysis

**DOI:** 10.1101/687467

**Authors:** Abate Dargie, Yossef Eshetie, Yared Asmare, Wendimeneh Shibabaw, Kefyalew Dagne

## Abstract

**Introduction:** maternal alcohol beverages consumption (any amount) during pregnancy can result in multiple major health and social problems both for the mother and fetus; including miscarriage, stillbirth, low birth weight, and prematurity. At the regional and national level, alcohol use prevalence data is a use full indicator for maternal and child health.

**Methods:** the researchers were searched for studies using a computerized search engine, main electronic databases, and other applicable sources. Observational studies (case-control, crosssectional and cohort) which assess the prevalence of alcohol use and associated factors among pregnant mothers in East Africa were eligible. Data was extracted thoroughly by two authors independently and screened for eligibility. The Pooled prevalence of alcohol use during pregnancy and its association with partner alcohol use was determined by using Epi data version 14 statistical software.

**Results:** the study included eighteen studies with the total sample size of 41,022 and The overall pooled prevalence of alcohol use during pregnancy from the random effects method was found to be 18.85% (95% CI; 11.26, 26.44). The overall weighted odds ration revealed that pregnant women partners’ alcohol use did not have a significant association with study subjects alcohol use during pregnancy; i.e. OR=**0.32** (**95% CI: −0.39, 1.03**).

**Conclusions:** The overall alcohol use (any amount) during pregnancy is higher in magnitude and pregnant mothers who had alcohol user partner had no association with their use of alcohol beverages. The prevalence of alcohol use during pregnancy may be underestimated in the current study due to social desirability bias. Since related study articles were found only in four East African countries, the region may be under-represented due to the limited number of studies included.

## 1. Introduction

Alcohol is the most commonly used teratogen and harmful substance that can cause a problem for pregnancy and places the mother at hazard but also unfavorable effects on the growing fetus which can permanent (1). According to the world health organization, there is no safe amount of alcohol, time of gestation and type of alcohol during pregnancy. Pregnant women never drink alcohol alone; when the mother drinks so does the fetus. The bad news here is the fetus cannot metabolize alcohol(2).

Perinatal alcohol use disorder can result in major organ birth flaws, neurodevelopmental disorders, Fetal alcohol syndromes and damage to multiple structures in the brain. Excessive alcohol consumption during pregnancy can result in lasting disabilities and numerous health and social problems for both mother and child, including miscarriage, the death of a fetus, low birth weight, and prematurity(3).The effect of excessive alcohol drinking on behavior, health, and society is a major public health problem globally (4). Furthermore, cognitive deficits have also been manifested in language, motor development, concentration, memory, and judgment. In addition, heart, muscle, kidney, vision, and hearing deficits have been substantiated the effect of alcohol use during pregnancy(5).

The disorderly drinking of alcohol in the first weeks of gestation may be associated with cases of spontaneous abortion, and its consumption between the third and eighth weeks of gestation might increase the risk of bodily malformations in the fetus (Fetal Alcohol Syndrome); which affects 33% of children born from mothers who consume more than 150 gram of alcohol per day. Moreover, children of women who drink alcohol moderately may present agitation, suction deficiency during feeding, irritability, sweating and abnormal sleep patterns, characterizing a condition of abstinence syndrome(6).

The prevalence of alcohol use (any amount) during pregnancy found to be higher in Africa countries including East Africa. From a compilation of the studies identified in Africa, the prevalence of alcohol use during pregnancy ranges from 2.5% to 37.9% (7–18). Different determinant factors are identified for maternal alcohol drinking and use disorder during pregnancy. Of them the mostly reported predictors are previous pregnancy obstetric complication, prior alcohol use, maternal mental health problem, partner alcohol use, making local brews, unplanned pregnancy and mothers educational level were frequently reported (7, 10, 12, 15, 16, 19–23). Local and standard Alcoholic beverages are rampant in East Africa and individuals drink them irrespective of gender and pregnancy status (2, 24–28). East African administrations are drafting different strategies currently to promote maternal and child health by giving special emphasis to pregnant women and the conceived fetus. Therefore the pooled prevalence data of maternal alcohol consumption (any amount) and its association with predictors is needed to evaluate the status of women alcohol use and the factors for their alcohol consumption.

## 2. Methods

### 2.1. Identification and study selection

The current study was conducted to quantify the pooled prevalence of alcohol use (any amount) and its association with newborn birth outcome. Articles were reviewed from national and international databases. The following databases were systematically searched: Boolean operator, Cochrane library, PubMed, EMBASE, Google Scholar, libraries and direct google search from conception to last January 2018. The reports will accessed using the following key terms/like Mesh terms; “alcohol use”, “alcohol consumption”, “alcohol drinking”, “pregnancy”, “anemia”, “preterm birth”, “low birth weight”, “adverse outcome”, and “each east Africa country”. The key terms were used individually and in combination through “AND” string. Each east African countries were combined with the above terms to intensively extract articles from the databases. Direct google search and library searching were done to identify grey literature. In addition, after the identification of studies and review articles, their lists of reference were searched to identify more eligible studies. The above database search strategy and terms were presented. This systematic review and meta-analysis used the PRISMA checklist to determine the eligibility of the articles included in the study.

### 2.2. Eligibility criteria

#### 2.2.1. Inclusion criteria

**Study area:** only studies done in East Africa countries were included

**Study design:** All observational studies (cross-sectional, cohort and case controls) that contain Original data; reporting the prevalence among alcohol use among pregnant women were considered.

**Language**: Literature published in the English language were included. **Population**: Studies conducted among pregnant women were considered

**Publication condition**: Both published and unpublished articles were considered. All published studies and grey literature from January first, from conception to January last, 2018 were considered for the review. Duplicate studies: here, the most comprehensive and/or recent study with the largest sample size was chosen during the abstraction.

#### 2.2.2. Exclusion criteria

The following studies were excluded from the review:

Studies which did not report the main outcome of the study were excluded. In addition, articles, which were not fully accessible after at least two-email contact with the corresponding author was excluded; because, in fact, it affects the quality of data in the absence of full text.

### 2.3. Data abstraction

Five reviewers (AD, YE, WS, YA, and KD) were screened the searched articles against the inclusion criteria through independent reading of the titles and abstracts, which were searched and accessed broadly. The full texts of these articles were accessed, and independent assessment was carried out by two reviewers; i.e. AD and YE, for eligibility based on the predetermined inclusion and exclusion criteria. Discrepancies between the reviewers were resolved through discussion and common consensus of all investigators. Multiple publications of the same study data from the included papers were extracted by YA; WS and KD independently. Assessment of the study quality was performed by four reviewers (AD, YE, YA, WS, and KD) independently and discrepancies between the evaluators were summarized with consensus. In the grading of the quality of studies, the reviewers were guided by the Newcastle– Ottawa Scale(29).

### 2.4. Outcome measurement

This systematic review and meta-analysis have two main outcomes. The primary outcome was to determine the weighted prevalence of alcohol use during pregnancy. The second outcome of the study was the association of partner alcohol use with pregnant women alcohol use. The prevalence was calculated by dividing the number of participants engaged alcohol use to the total number of participants who have been included in the study (sample size) multiplied by 100. Regarding the predictor variable, we calculated the odds ratio from the primary studies using their reported odds ratio and its confidence interval.

### 2.5. Quality assessment

The abilities of each of the finding reports in the systematic review were assessed by using a checklist adjusted from the Joanna Briggs Institute (JBI) Critical Appraisal for Study Papers(30).

To assess the quality of studies, the researchers used the tool Newcastle-Ottawa Scale adapted for cross-sectional studies quality assessment. The tool has indicators consisting of three main sections; the first section has five stars and assesses the methodological quality of each study. The second section of the tool evaluates the comparability of the studies. The last part of the tool measures the quality of the original articles with respect to their statistical analysis(29). Using the tool as a protocol, the two authors independently evaluated the qualities of the original articles. The quality of the studies was evaluated by using these indicators; those with medium (fulfilling 50% of quality assessment criteria) and high quality (6 out of 10 scales) were included for analysis. By taking the mean score of the two researchers, disagreements of their assessment results were resolved (see table 1).

**Table 1:** Descriptive summary of 18 studies included in the meta-analysis of the prevalence of alcohol use during pregnancy in East Africa, 2019.

| Author | Publication Year | Country | Total Samples | Response Rate (%) | Prevalence (95% CI) | Quality score (10 pt.) |
| --- | --- | --- | --- | --- | --- | --- |
| Isaksen et al.(12) | 2015 | Tanzania | 34090 | 100 | 34.1(33.6,34.6) | 8 |
| Popova et al.(7) | 2016 | Uganda | 505 | 100 | 16(12.8,19.2) | 7 |
| Anteab et al.(22) | 2014 | Ethiopia | 810 | 100 | 34(30.74, 37.26) | 7 |
| Namagembe et al.(16) | 2010 | Uganda | 618 | 98.7 | 25(21.56,28.44) | 7 |
| Mpelo et al.(15) | 2018 | Tanzania | 365 | 100 | 15.1(11.43,18.77) | 6 |
| Tesso et al.(10) | 2017 | Ethiopia | 296 | 99 | 37.9(32.37,43.43) | 6 |
| Abate et al.(19) | 2018 | Ethiopia | 685 | 100 | 16.1(13.35,18.85) | 7 |
| Obse et al.(17) | 2013 | Ethiopia | 296 | 98.9 | 37.9(32.35,43.45) | 6 |
| Demelash et al.(33) | 2015 | Ethiopia | 136 | 94.9 | 24.8(17.35,32.25) | 6 |
| Hanlon et al.(11) | 2009 | Ethiopia | 515 | 100 | 3.3(.76,4.84) | 7 |
| Mosha and Philemon(14) | 2010 | Tanzania | 157 | 100 | 2.5(0.06,4.94) | 6 |
| Namagembe et al.(16) | 2010 | Uganda | 618 | 98.7 | 30(26.36,33.64) | 7 |
| Aboye et al.(20) | 2018 | Ethiopia | 308 | 100 | 16.2(12.09,20.31) | 6 |
| Wagura et al.(18) | 2018 | Kenya | 322 | 100 | 6.5(3.41,9.19) | 6 |
| Demelash et al.(33) | 2015 | Ethiopia | 408 | 94 | 18.8(14.86,22.76) | 7 |
| Asmare et al.(32) | 2018 | Ethiopia | 453 | 94.7 | 11.4(8.16,14.64) | 7 |
| Ahmed et al.(21) | 2018 | Ethiopia | 286 | 97.6 | 7.2(4.17,10.23) | 6 |
| Makayoto et al.(13) | 2013 | Kenya | 300 | 100 | 3.7(1.56,5.84) | 6 |

### 2.6. Statistical analysis

Data were extracted from each appropriate article using a Microsoft Excel spreadsheet software and imported into STATA Version 14 software for analysis. Stata version 14 was used to calculate the pooled effect size with 95% confidence intervals of alcohol use. Statistical heterogeneity between the studies and publication bias were performed with appropriate test statistics. A weighted inverse variance random-effects model was used to estimate the pooled prevalence of alcohol use during pregnancy in the current meta-analysis. The included eighteen studies outcome were evaluated for heterogeneity and publication bias. Heterogeneity across the studies was assessed using I^2^ statistic were 25, 50 and 75% representing low, moderate and high heterogeneity respectively(31).A Funnel plot and Egger’s regression test were used to check publication bias(15). Consequently, the analysis showed a substantial heterogeneity of egger test (p < 0.001) and I^2^ statistics (I^2^ = 99.5%). The funnel plot for publication bias also showed no symmetry verifying the presence of bias. Forest plot was also used to show the pooled prevalence of alcohol use during pregnancy. In addition, to minimize the random variations between the point-estimates of the primary study, subgroup analysis was done based on the country, study design and sample size.

## 3. Results

### 3.2. Search Results

All studies were done in East Africa countries and published in an indexed journal and grey literature from direct google search and libraries searching were eligible for this study. A total of eighteen studies were included and fifteen of them were cross-sectional studies, one cohort and the remaining two were case-control studies. The advanced search by using PUBMED/EMBASE Database 29,749 articles, direct google search 12,963 articles, and library two articles have resulted. After immense efforts of reviewing the accessed articles, eighteen eligible research articles were included for the quantitative meta-analysis with a total sample size of 40,722 pregnant mothers who consume alcohol during pregnancy (see Fig. 1).

**Fig. 1:**
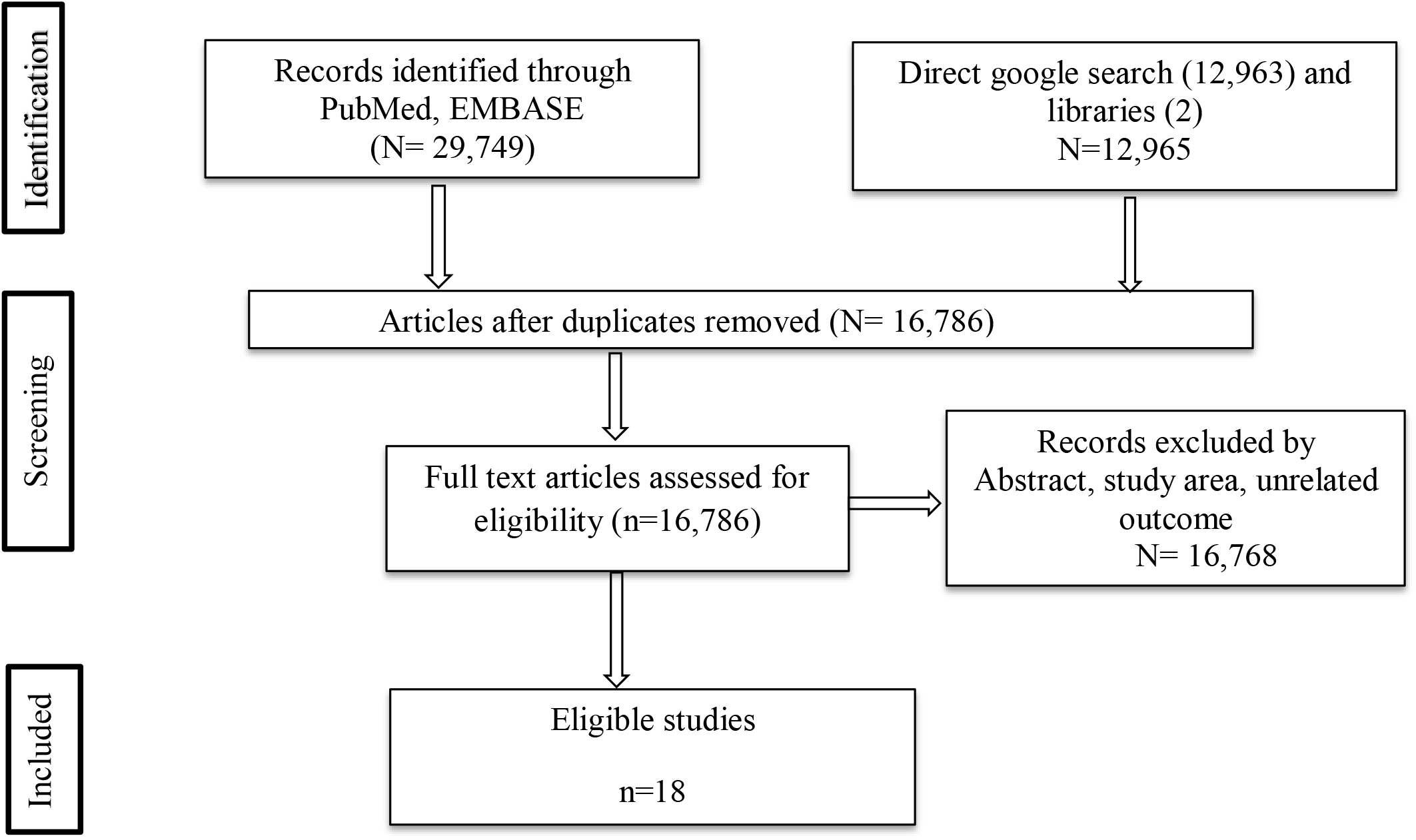
Selection Process for the Studies Included in the Systematic Review and Meta-Analysis in East Africa, 2019.

Information about authors, publication year, population, study area, study design, outcome and main results from the selected articles were extracted and summarized on the table. The overall response rate was between nighty four percent (94%) to a hundred percent (100%).

The current study was conducted by compiling studies done in different east African countries; Ethiopia (10, 11, 17, 19–22, 32, 33), Uganda(7, 16), Tanzania(12, 14, 15), and Kenya(13, 18). Fifteen cross-sectional studies and two case-control and one cohort study articles were used for the final analysis. The sample size of the studies ranging from 136 to 34,090 pregnant women.

### 3.2. The pooled prevalence of alcohol use during pregnancy

In the current systematic review and meta-analysis, the pooled prevalence estimates of maternal alcohol use during pregnancy were showed by forest plot (Fig. 2). In the random effect model, weight for every study was given based on individual study effect size and sample size. In this case, the weight given for each included studies is presented in figure 3. In this review, the prevalence of each study ranged from 2.5 % to 37.9% with substantial heterogeneity across studies (I^2^ = 99.5%). The overall pooled prevalence of alcohol use during pregnancy from the random effects model was found to be 18.85% (95% CI; 11.26, 26.44).

**Figure 2:**
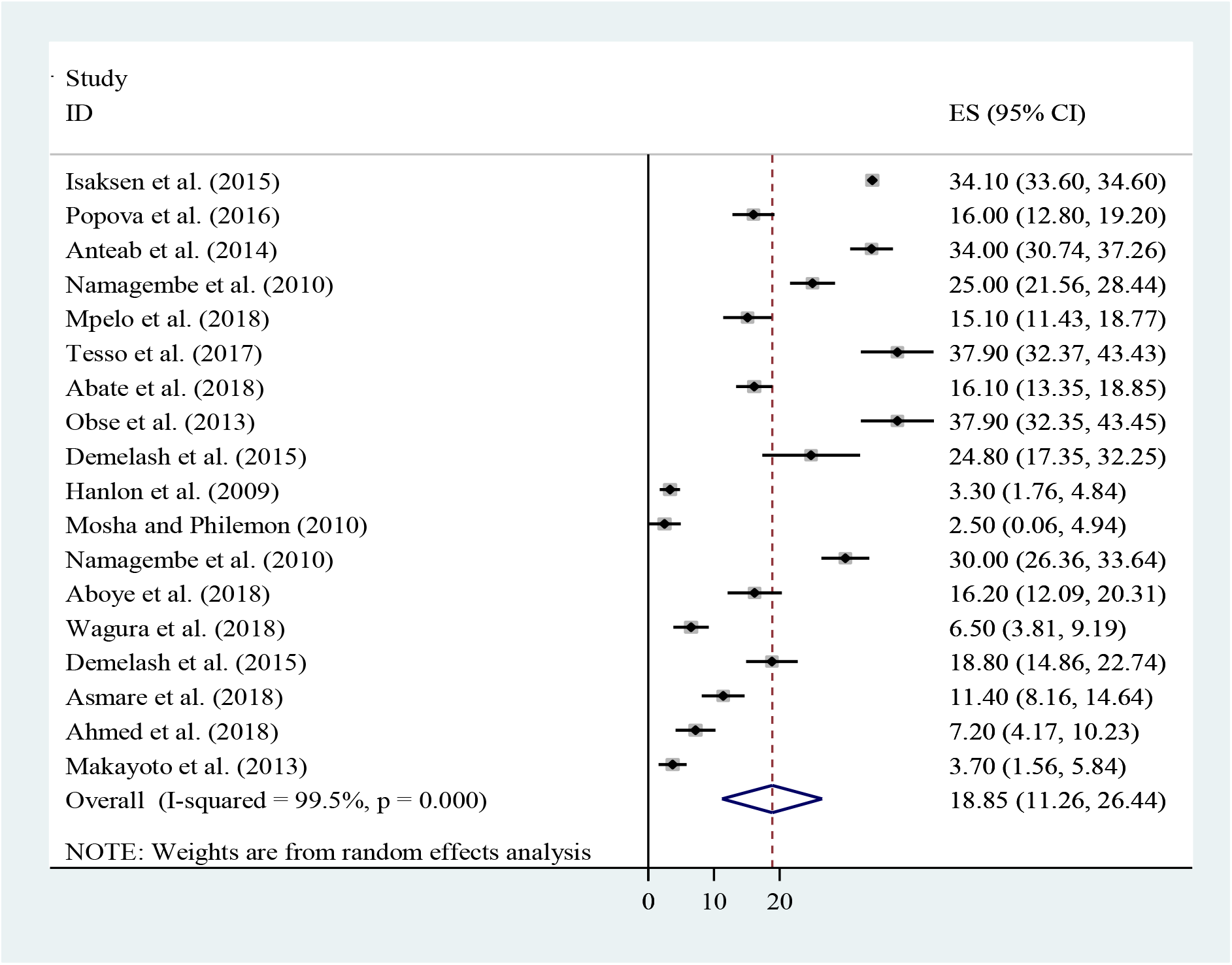
Forest plot of the overall pooled prevalence *of prenatal alcohol exposure as measured by maternal self-reports in East Africa, 2019*

**Fig. 3:**
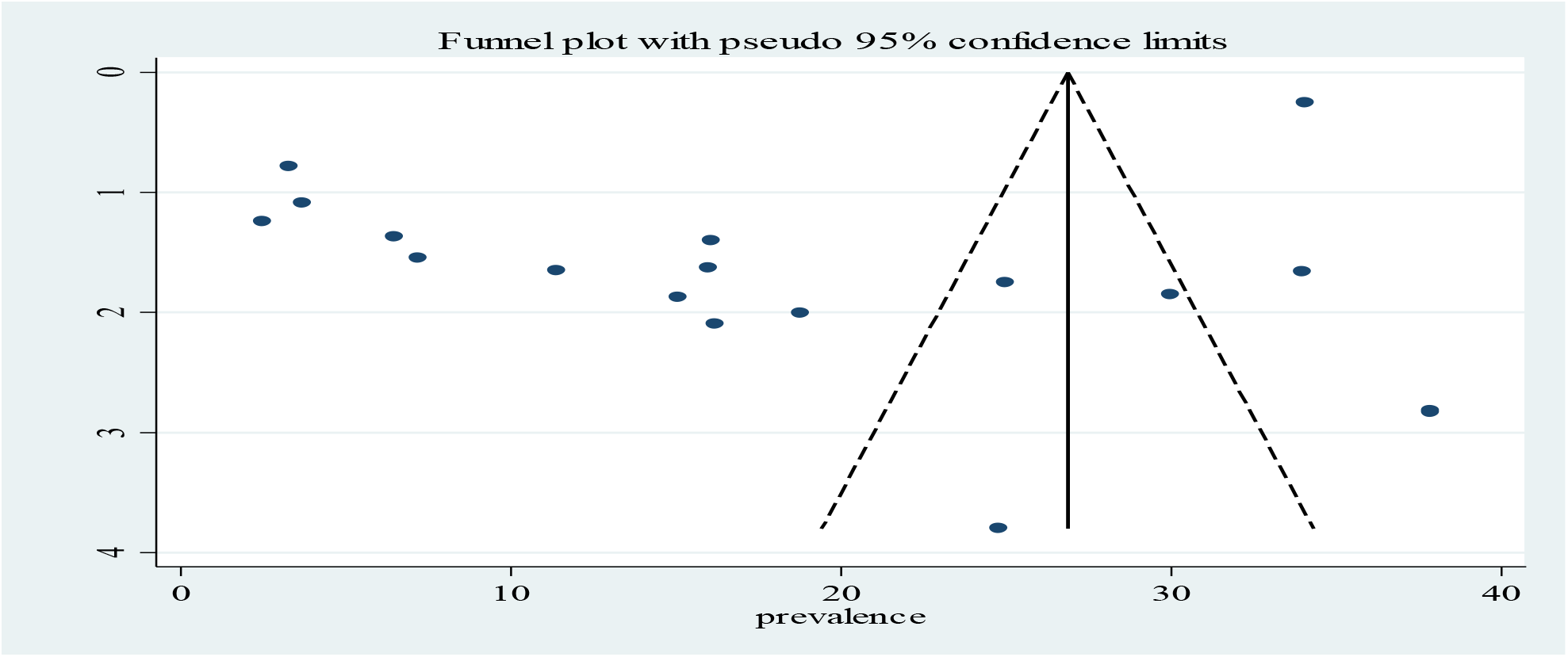
Funnel plot of the prevalence of prenatal alcohol exposure as measured by maternal selfreports in East Africa, 2019

**Fig. 3.**
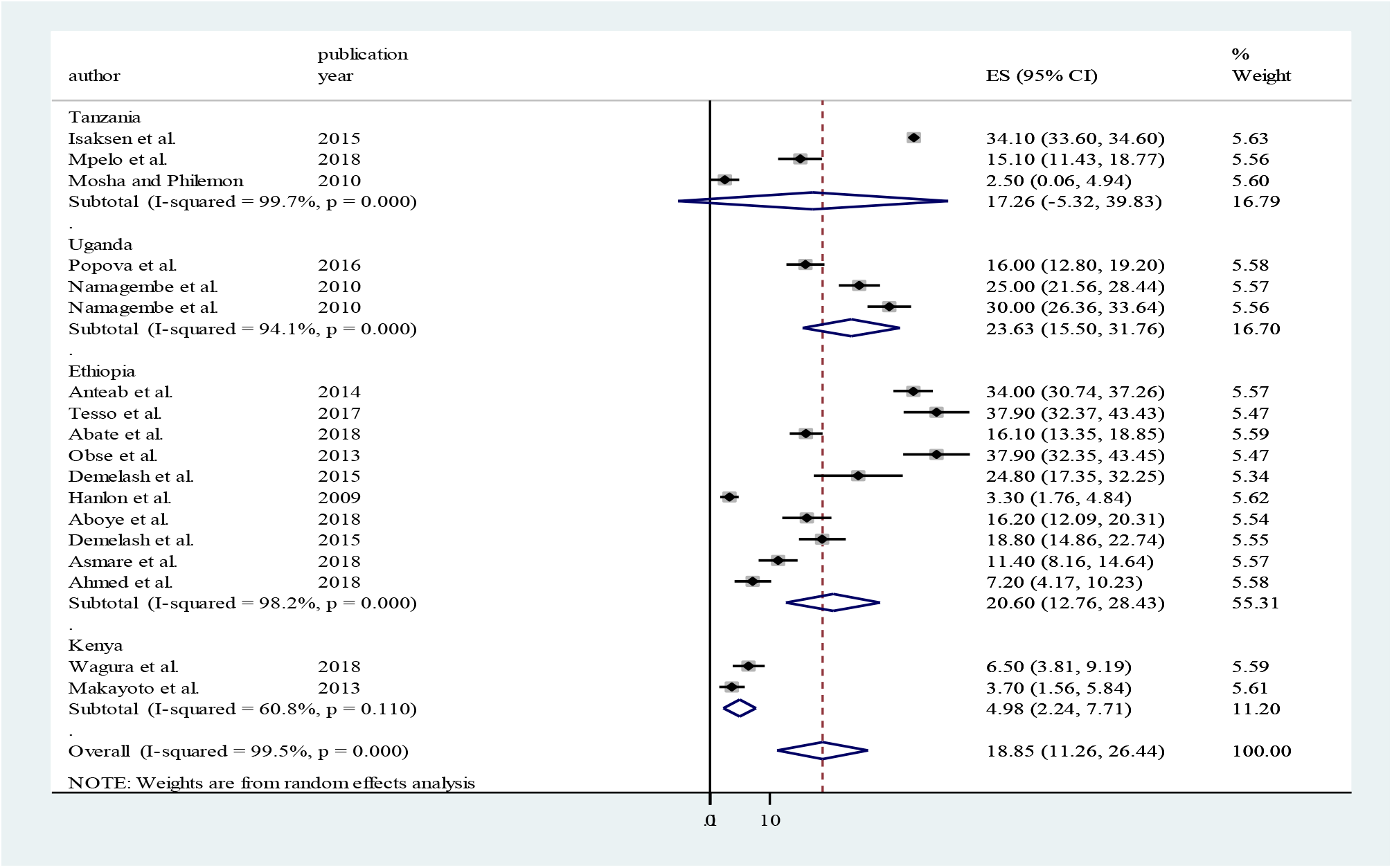
The subgroup prevalence of alcohol use during pregnancy by country in East Africa, 2019

On eyeball test, the funnel plot was found to be asymmetric and Egger’s test of the intercept (Bo) was found to be significant at a p-value of 0.00 which confirms that there is a publication bias within articles which needs subgroup analysis by different characteristics of the included journals.

### 3.3. Subgroup analysis of the research articles

Subgroup analysis was carried out based on the country, study design and number of samples included (see fig). Accordingly, the highest prevalence was reported in Uganda, 23.63 % (95%CI: 15.5, 31.76) followed by Ethiopia, 20.6% (95%CI: 12.76, 28.43). In the current systematic review and meta-analysis, the lowest prevalence was recorded in cross-sectional studies, 21.59% (95%CI: 14.01, 29.18). The single cohort study done in Ethiopia showed a lower magnitude of alcohol use during pregnancy. In addition, the reviewers performed subgroup analysis based on the number of samples included in each the study; the largest magnitude was found in studies done with a sample size of five hundred and above;22.63% (95%CI:10.67, 34.59), (Table 2).

**Fig. 4.**
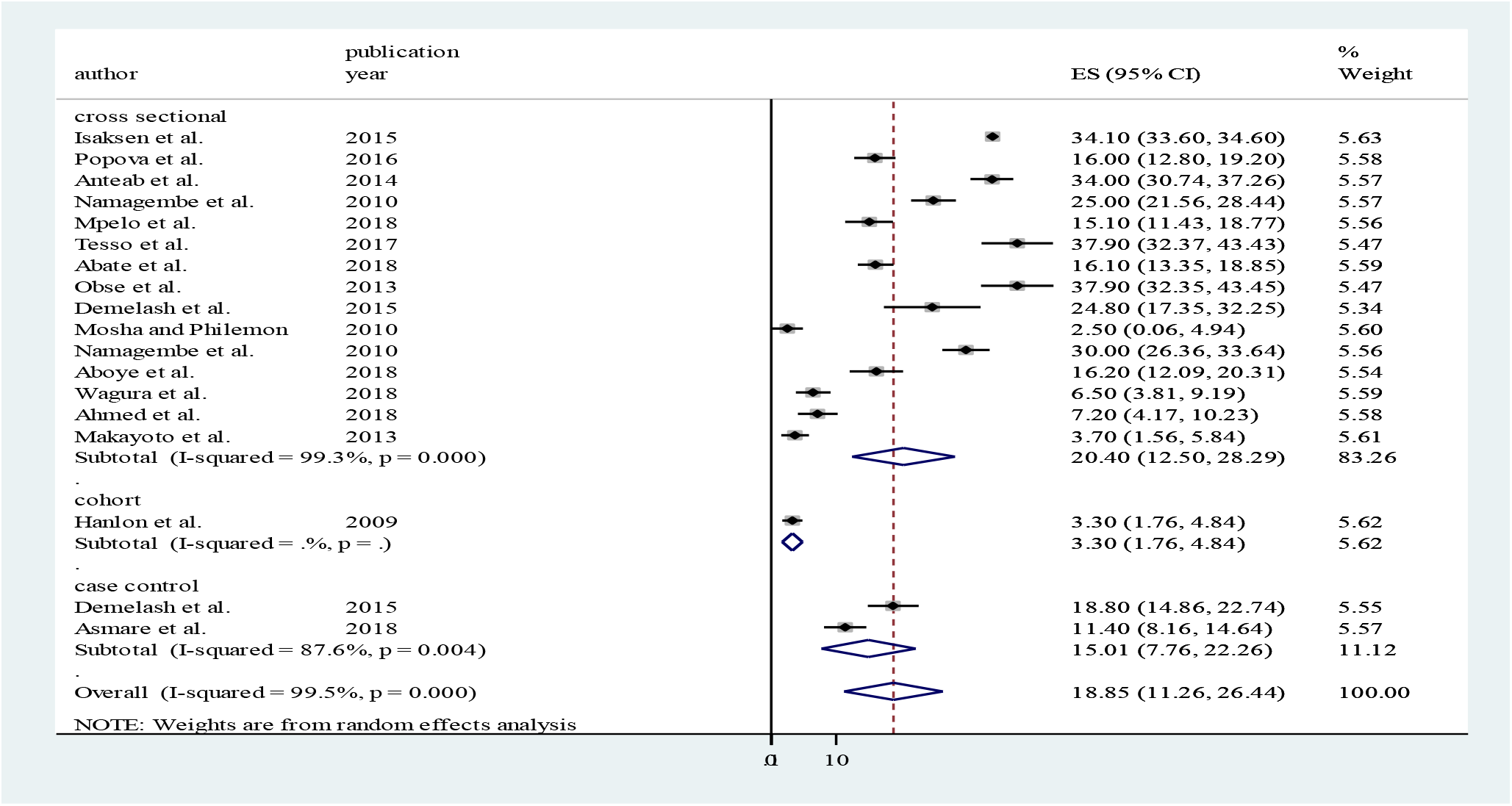
The subgroup prevalence of alcohol use during pregnancy by study design in East Africa, 2019

**Fig. 5.**
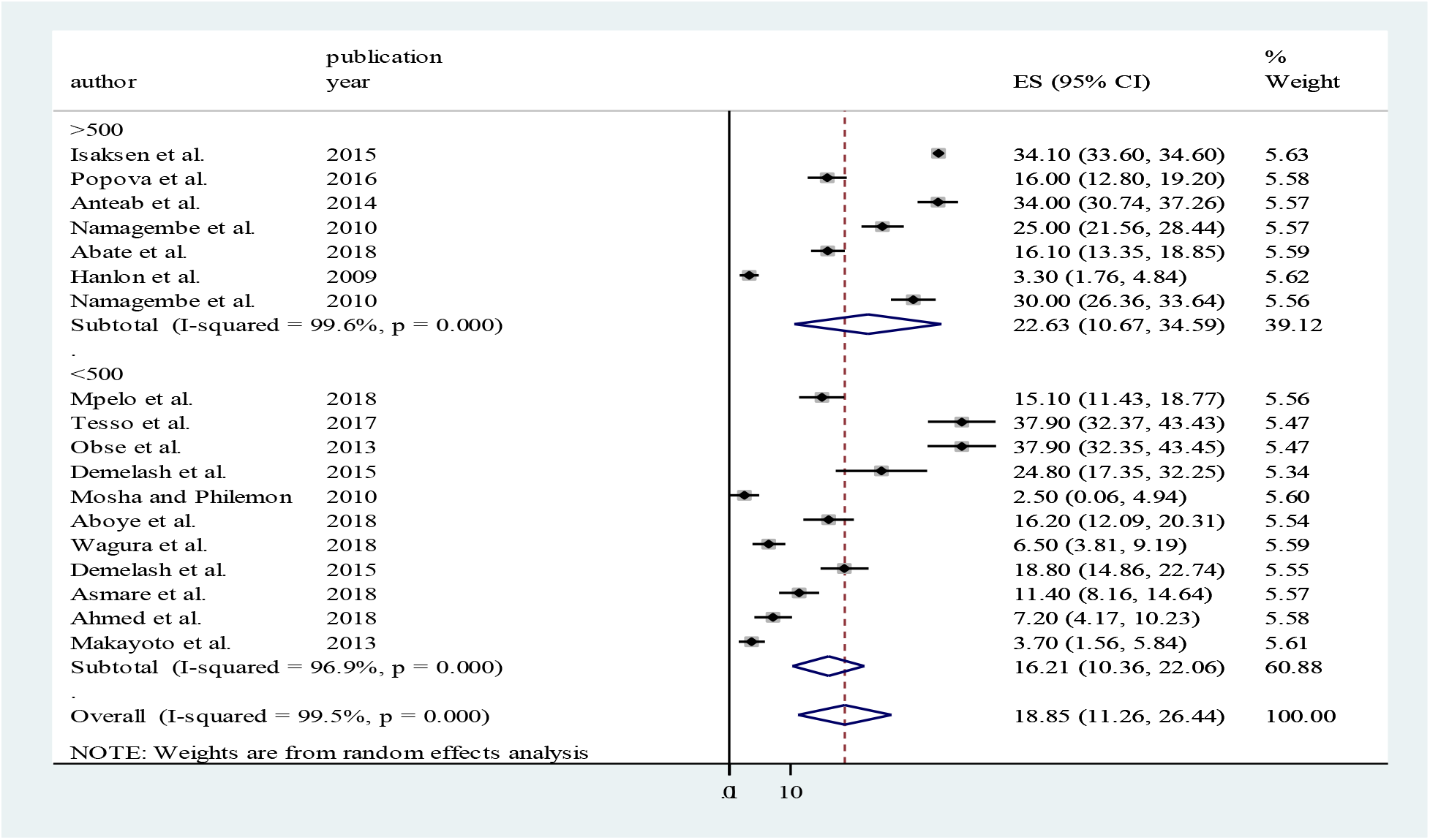
The subgroup prevalence of alcohol use during pregnancy by sample size included in the study in East Africa, 2019

**Table 2:** Summary of the subgroup analysis of the *prevalence of prenatal alcohol exposure as measured by maternal self-reports in East Africa, 2019*

| Subgroups | Characteristics | Included studies | Sample size | Prevalence(95% CI) |
| --- | --- | --- | --- | --- |
| **By country** | Tanzania | 3 | 34,612 | 17.3(-5.32,39.83) |
|  | Uganda | 3 | 1,725 | 23.63 (15.5,31.76) |
|  | Ethiopia | 10 | 4,063 | 20.6( 12.76,28.43) |
|  | Kenya | 2 | 622 | 4.98 (2.24,7.71) |
| **By design** | Cohort | 1 | 515 | 3.3 (1.76,4.84) |
|  | Cross-sectional | 14 | 39,459 | 21.59 (14.01,29.18) |
|  | Case control | 2 | 748 | 15.01(7.76,22.26) |
| **By included sample size** | ≥ 500 | 7 | 37,825 | 22.63 (10.67, 34.59) |
|  | <500 | 11 | 2,897 | 16.21(10.36,22.06) |
| **Over all** |  | **17** | **41,022** | **18.85%( 11.26,26.44)** |

### 3.4. Association of partner alcohol use with alcohol use during pregnancy

Is having an alcoholic partner a risk factor for pregnant women alcohol use? In five studies, having an alcoholic partner had no significant association with alcohol use during pregnancy when compared to counterparts (Fig. 7). Five studies (13, 15–17, 22), which assessed the association of partners alcohol use with maternal alcohol use during pregnancy was included to examine the association. Eyeball examination of the funnel plot also showed no variability across the individual studies (fig. 6).

**Fig. 6:**
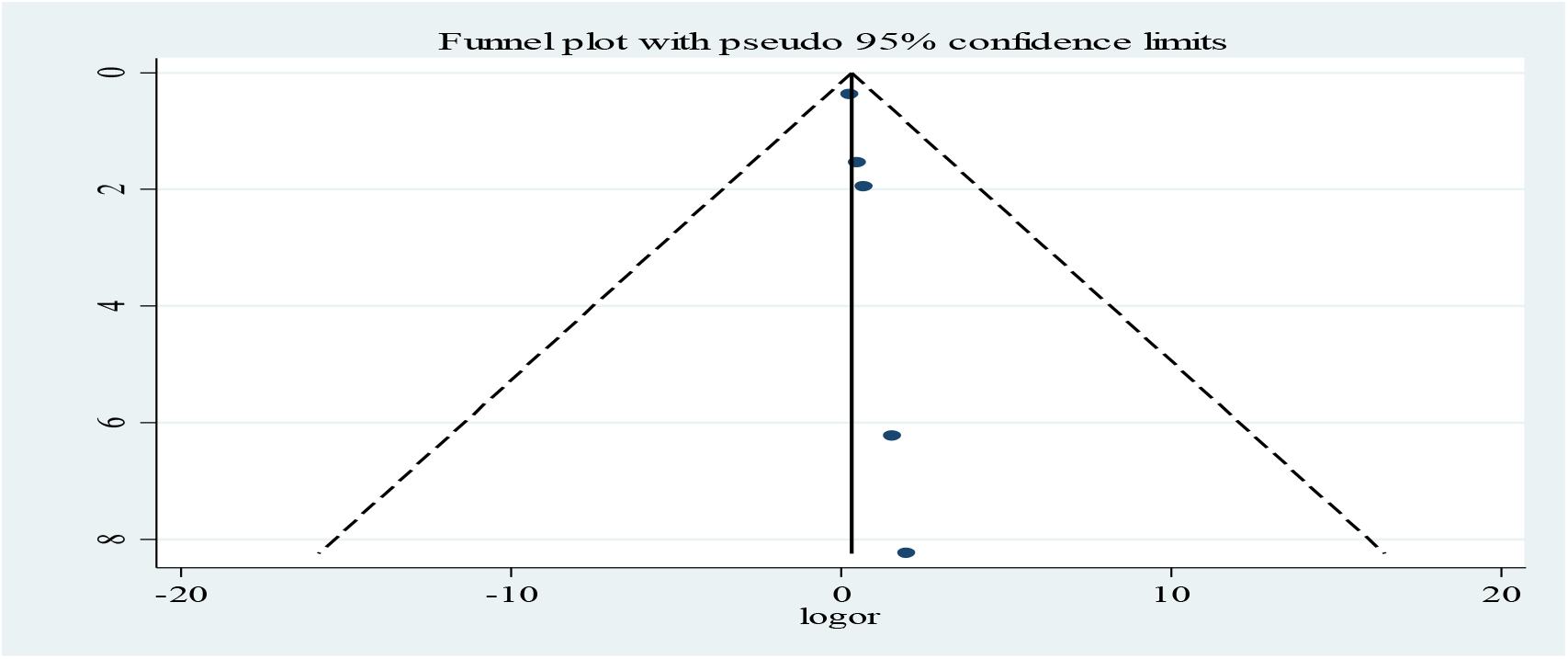
Shows funnel plot of the parent’s alcohol use to show the statistical heterogeneity in East Africa, 2019

The pooled odds ratio was **0.32** (**95% CI: −0.39, 1.03**), **I**^2^= 0.0% with a p-value of 0.997); the pooled odds ration showed that there was no significant association between partner alcohol use and mothers’ alcohol use during pregnancy (fig.7). Heterogeneity chi-square= 0.15 (d.f. =4), P-value=0.997 tests reveal no publication bias.

**Fig. 7.**
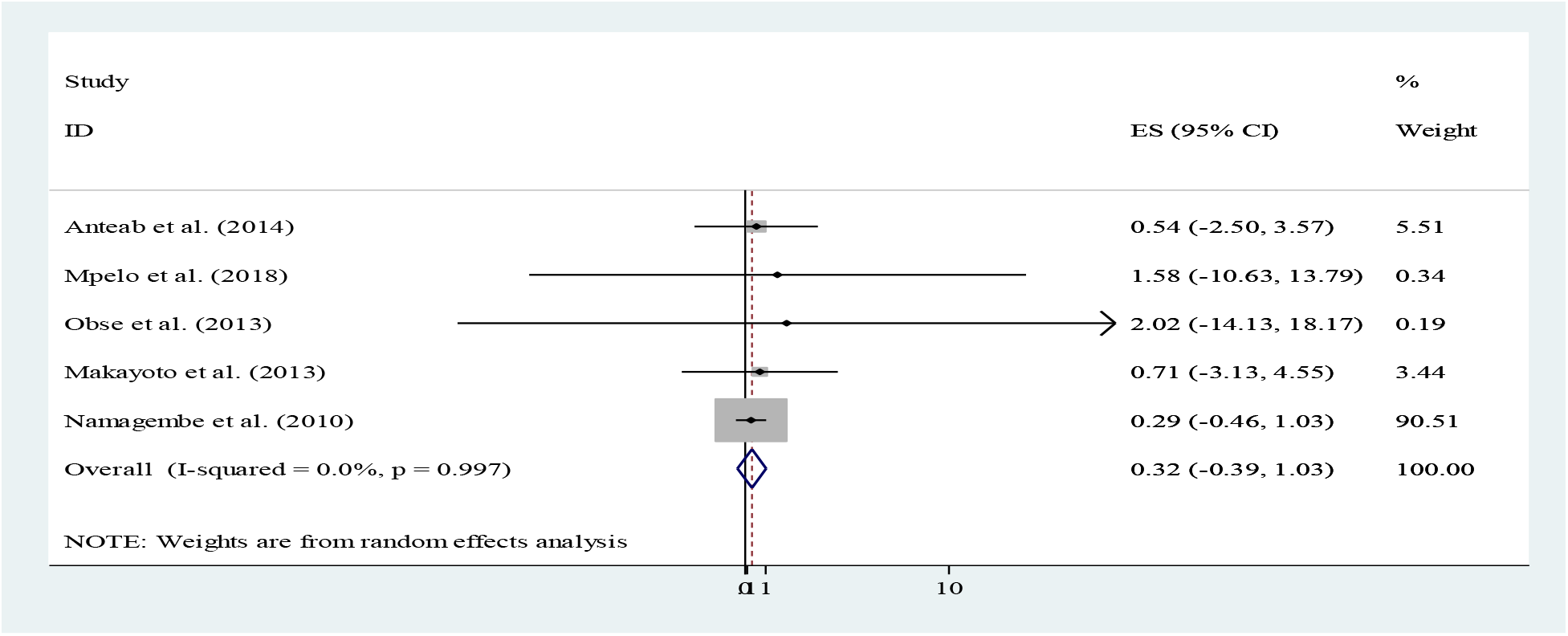
The pooled odds ratio of alcohol use during pregnancy in East Africa, 2019

## 4. Discussions

The results of this study reveal that in east African countries (Ethiopia, Tanzania, Uganda, and Kenya), the weighted prevalence of alcohol use among pregnant women is high; 18.85 %(95%CI= 11.26, 26.44). The prevalence of alcohol use during pregnancy varied within countries. Between-country variations in the prevalence of alcohol use during pregnancy likely originate not only from variation in maternal drinking behaviors but also from political, religious, ideological and cultural differences in East Africa. In East Africa, varying pooled weight of alcohol use have been found; the highest reported prevalence of alcohol use found in Uganda (23.6%); followed by Ethiopia (20.6%), Tanzania (17.3%) and Kenya (5%); which is clearly high rates of alcohol drinking during pregnancy (see fig.3).

Alcohol consumption during pregnancy may be influenced by lack of access to health information regarding the negative outcomes of alcohol consumption during pregnancy. In a study among pregnant women in Uganda, it was reported that 53.5% of respondents were not believed that alcohol consumed during pregnancy affects baby’s health and only 5 % of respondents had never attended formal education. Almost 61%(n=233) of pregnant mothers were not informed about the effect of alcohol drinking during pregnancy among Ethiopian women and 26%(n=99) study subjects unable to read and write (16, 19).

Easily availability of alcohol also influences consumption patterns. In many countries of the East African Region, alcohol is widely available due to either weak licensing systems or execution procedures. For instance, pubs and bars working on a 24-hour basis and low-cost alcoholic beverages are commonly retailed without a license; are very common in Ethiopia, Uganda, Kenya, and Tanzania. Policies to regulate alcohol publicizing in almost all African countries are also very fragile(26).

The unsatisfactory rules concerning alcohol affect drinking in general. Integral cultural practices may also vary the prevalence at which alcohol is consumed during pregnancy. For instance, in Africa, there are noticeable variances in the cultural practices witnessed between regions, countries and communities and their beliefs surrounding alcohol consumption in general. (34)

The meta-analyses on the consumption of any amount of alcohol consumed during pregnancy for both Tanzania and Uganda were based on only three studies; and only two studies in Kenya. The actual prevalence value of alcohol use during pregnancy may fluctuate from the true prevalence due to most East Africa countries were not included in the current study (no related articles found) and also limited research articles were found in the included countries.

Yet, our study offers an operational estimate of the prevalence of alcohol use during pregnancy in countries that do not currently have actual data. The current prevalence estimates (meta-analysis or regression estimations) are useful indicators of the communal health problem of alcohol drinking during pregnancy and provide a basis for resource allocation for prevention strategies. The results of this study will optimistically encourage countries to conduct their own studies in order to estimate their own prevalence statistics on alcohol drinking during pregnancy.

## 5. Conclusions

The overall alcohol use (any amount) during pregnancy is higher in magnitude and pregnant mothers who had alcohol user partner had no association with their use of alcohol beverages.Since related study articles were found only in four East African countries, the region may be under-represented due to the limited number of studies included.

## 6. Recommendations

Africa is a highly varied continent in socioeconomic status, religion, and ethnicity and all recommendation provided below will not be applicable to all countries. Though, the current study findings highlight that there is a need to educate all pregnant women about the possible harmful effects of alcohol exposure on the growing fetus and to establish screening protocol and provide brief interventions, where appropriate, to all pregnant women at the health service institutions and community level. Additionally, both the availability and the effectiveness of substance use disorder treatment programs for women of reproductive age and the mothers of children with fetal alcohol syndrome should be expanded and strengthened.

## 7. The strength of the study

Strengths of this study include: - having intensive searching strategy along with demanding inclusion and exclusion criteria, and statistical analyses.

## 8. Limitation of the study

The self-reported data on alcohol consumption during pregnancy are sensitive to social desirability bias and recall bias. Therefore, the prevalence of alcohol use during pregnancy may be underestimated in the current study. Only English articles were considered to be included in this review. In addition, the majority of the studies included in this review were cross-sectional in nature as a result; the outcome variable might be affected by other confounding variables.

## 9. Declarations

### 9.1. Availability of data

Data will be available upon request of the corresponding author.

### 9.2. Authors’ contributions: AD

Conception of the research protocol, study design, literature review, data extraction, data analysis, interpretation and drafting the manuscript. YA, WS, and YE: data extraction, quality assessment, data analysis and reviewing the manuscript. All authors have read and approved the manuscript.

### 9.3. Ethics approval and consent to participate

Since the review was concerned with the published research articles, there was no need for ethical approval and/or additional consent from participants.

### 9.4. Funding and Consent for publication

not applicable

### 9.5. Competing interest

The authors declare that they have no competing interests.

## 9.6. Acknowledgments

it is my pleasure to acknowledge all the reviewers for their time devotion and commitment during the process of the research. Finally, we would like to appreciate African Mental Health Research Initiative (AMARI) which is funded through the DELTAS Africa initiative (DEL-15-01) for supporting the co-author (Kefyalew Dagne) financially as well as through providing various capacity building initiatives including systematic review and meta-analysis writing workshop which facilitated the writing of the present manuscript.

## References

1. Manchikanti L. National drug control policy and prescription drug abuse: facts and fallacies. Pain physician. 2007;10(3):399.

2. Department WHOSA, Health WHODoM, Abuse S, Organization WH. Global status report: alcohol policy: World Health Organization; 2004.

3. WHO Guidelines for the identification and management of substance use and substance use disorders in pregnancy. WHO Library Cataloguing-in-Publication Data. 2014.

4. World Health Organization Geneva S. Global Status Report on Alcohol and Health. 2011.

5. Green JH. Fetal Alcohol Spectrum Disorders: Understanding the Effects of Prenatal Alcohol Exposure and Supporting Students. Journal of School Health. 2007;77:103–8.

6. Oliveira TR, SMF S. O consumo de bebida alcoólica pelas gestantes:um estudo exploratório. 2007;11(4):632–8.

7. Popova S, Lange S, Probst C, Shield K, Kraicer-Melamed H, Ferreira-Borges C, et al. Actual and predicted prevalence of alcohol consumption during pregnancy in the WHO African Region. Tropical Medicine & International Health. 2016;21(10):1209–39.

8. Adusi-Poku Y, Edusei AK, Bonney AA, Tagbor H, Nakua E, Otupiri E. Pregnant women and alcohol use in the Bosomtwe district of the Ashanti region-Ghana. African journal of reproductive health. 2012;16(1).

9. Lange S, Shield K, Koren G, Rehm J, Popova S. A comparison of the prevalence of prenatal alcohol exposure obtained via maternal self-reports versus meconium testing: a systematic literature review and meta-analysis. BMC pregnancy and childbirth. 2014;14(1):127.

10. Fekadu Yadassa Tesso* LAW, and Dagmawit Birhanu Kebede. Magnitude of Substance Use and Associated Factors among Pregnant Women Attending Jimma Town Public Health Facilities, Jimma Zone, Oromia Regional State Southwest Ethiopia Clinics in Mother and Child Health-Omics international. 2017;14(4):1–5.

11. Hanlon C, Medhin G, Alem A, Tesfaye F, Lakew Z, Worku B, et al. Impact of antenatal common mental disorders upon perinatal outcomes in Ethiopia: the P-MaMiE population-based cohort study. Tropical Medicine & International Health. 2009;14(2):156–66.

12. Isaksen AB, Østbye T, Mmbaga BT, Daltveit AK. Alcohol consumption among pregnant women in Northern Tanzania 2000–2010: a registry-based study. BMC pregnancy and childbirth. 2015;15(1):205.

13. Makayoto LA, Omolo J, Kamweya AM, Harder VS, Mutai J. Prevalence and associated factors of intimate partner violence among pregnant women attending Kisumu District Hospital, Kenya. Maternal and child health journal. 2013;17(3):441–7.

14. Mosha TC, Philemon N. Factors influencing pregnancy outcomes in Morogoro Municipality, Tanzania. Tanzania journal of health research. 2010;12(4):243–51.

15. Mpelo M, Kibusi SM, Moshi F, Nyundo A, Ntwenya JE, Mpondo BC. Prevalence and Factors Influencing Alcohol Use in Pregnancy among Women Attending Antenatal Care in Dodoma Region, Tanzania: A Cross-Sectional Study. Journal of pregnancy. 2018;2018.

16. Namagembe I, Jackson LW, Zullo MD, Frank SH, Byamugisha JK, Sethi AK. Consumption of alcoholic beverages among pregnant urban Ugandan women. Maternal and child health journal. 2010;14(4):492–500.

17. Obse N, Mossie A, Gobena T. Magnitude of anemia and associated risk factors among pregnant women attending antenatal care in Shalla Woreda, West Arsi Zone, Oromia Region, Ethiopia. Ethiopian journal of health sciences. 2013;23(2):165–73.

18. Wagura P, Wasunna A, Laving A, Wamalwa D. Prevalence and factors associated with preterm birth at kenyatta national hospital. BMC pregnancy and childbirth. 2018;18(1):107.

19. Abate Dargie KD, Surafel Habte. prevalence of problematic alcohol use and associated factors among pregnant women, North Shoa, Debre Berhan, Ethiopia, 2018. BMC pregnancy and childbirth. 2018;18.

20. Aboye W, Berhe T, Birhane T, Gerensea H. Prevalence and associated factors of low birth weight in Axum town, Tigray, North Ethiopia. BMC research notes. 2018;11(1):684.

21. Ahmed S, Hassen K, Wakayo T. A health facility based case-control study on determinants of low birth weight in Dassie town, Northeast Ethiopia: the role of nutritional factors. Nutrition journal. 2018;17(1):103.

22. Anteab K, Demtsu B, Megra M. Assessment of Prevalence and Associated Factors of Alcohol Use during Pregnancy among the dwellers of Bahir-Dar City, Northwest Ethiopia, 2014. 2014.

23. Birhanu AM, Bisetegn TA, Woldeyohannes SM. High prevalence of substance use and associated factors among high school adolescents in Woreta Town, Northwest Ethiopia: multi-domain factor analysis. BMC public health. 2014;14(1):1186.

24. Ashenafi M. A review on the microbiology of indigenous fermented foods and beverages of Ethiopia. Ethiopian Journal of Biological Sciences. 2006;5(2):189–245.

25. Ashenafi M, Mehari T. Some microbiological and nutritional properties of Borde and Shamita, traditional Ethiopian fermented beverages. Ethiopian Journal of Health Development. 1995;9:105–10.

26. Drivdal L, Lawhon M. Plural regulation of shebeens (informal drinking places). South African Geographical Journal. 2014;96(1):97–112.

27. Nikander P, Seppälä T, Kilonzo G, Huttunen P, Saarinen L, Kilima E, et al. Ingredients and contaminants of traditional alcoholic beverages in Tanzania. Transactions of the Royal Society of Tropical Medicine and Hygiene. 1991;85(1):133–5.

28. Teramoto Y, Sato R, Ueda S. Characteristics of fermentation yeast isolated from traditional Ethiopian honey wine, ogol. African Journal of Biotechnology. 2005;4(2):160–3.

29. Stang A. Critical evaluation of the Newcastle-Ottawa scale for the assessment of the quality of nonrandomized studies in meta-analyses. European journal of epidemiology. 2010;25(9):603–5.

30. Zeng X, Zhang Y, Kwong JS, Zhang C, Li S, Sun F, et al. The methodological quality assessment tools for preclinical and clinical studies, systematic review and meta-analysis, and clinical practice guideline: a systematic review. Journal of evidence-based medicine. 2015;8(1):2–10.

31. Higgins JP, Thompson SG, Deeks JJ, Altman DG. Measuring inconsistency in meta-analyses. Bmj. 2003;327(7414):557–60.

32. Asmare G, Berhan N, Berhanu M, Alebel A. Determinants of low birth weight among neonates born in Amhara Regional State Referral Hospitals of Ethiopia: unmatched case control study. BMC research notes. 2018;11(1):447.

33. Demelash H, Motbainor A, Nigatu D, Gashaw K, Melese A. Risk factors for low birth weight in Bale zone hospitals, South-East Ethiopia: a case–control study. BMC pregnancy and childbirth. 2015;15(1):264.

34. Kaba AJ. The Spread of Christianity and Islam in Africa: A Survey and Analysis of the Numbers and Percentages of Christians, Muslims and Those Who Practice Indigenous Religions. Western Journal of Black Studies. 2005;29(2).

